# XBP1s as a Therapeutic Target to Preserve Retinal Function During Aging and Neurodegeneration

**DOI:** 10.1101/2025.07.03.662890

**Authors:** Leonel E. Medina, Joel Barría, Max Chacón, Maximiliano Orellana, Daniela Ponce, David Neira, Francisco Miqueles, Claudia Duran-Aniotz, Joaquín Araya-Arriagada, Bruno Cerda, Claudio Hetz, Cristóbal Rojas, Adrian G. Palacios

**Affiliations:** Departamento de Ingeniería Biomédica, Universidad de Santiago de Chile, Santiago, Chile; Facultad de Medicina, Universidad Diego Portales, Santiago, Chile; Departamento de Ingeniería Informática, Universidad de Santiago de Chile, Santiago, Chile; Instituto de Neurociencia and Centro Interdisciplinario de Neurociencia de Valparaíso, Facultad de Ciencias, Universidad de Valparaíso; Latin American Institute for Brain Health, Universidad Adolfo Ibanez, Santiago, Chile; Escuela de Tecnología Médica, Facultad de Salud, Universidad Santo Tomás, Santiago, Chile; Instituto de Ingeniería Matemática y Computacional, Pontificia Universidad Católica de Chile, Santiago, Chile; Centro Nacional de Inteligencia Artificial, Chile; Institute of Biomedical Sciences, Universidad de Chile, Santiago, Chile; Instituto de Sistemas Complejos de Valparaíso, Valparaíso, Chile

## Abstract

Loss of physiological complexity, characterized by reduced adaptive multiscale coordination among system components, is increasingly recognized as a hallmark of aging and neurodegenerative disease. The retina, a window into the brain, offers a unique, accessible platform to monitor neurodegenerative disorders such as Alzheimer’s disease (AD). Here, we investigate the therapeutic potential of the unfolded protein response transcription factor XBP1s in preserving retinal function during aging and AD-related pathology in murine models. Using micro-electroretinography with multielectrode arrays, we recorded retinal responses to chirp and white noise stimuli in four mouse models: wild-type (WT), XBP1s-overexpressing (Tg^XBP1s^), AD model (5xFAD), and their crossbreed (Tg^XBP1s^/5xFAD) at approximately 3 and 7 months of age. We assessed retinal signals through entropy-based complexity measures and wavelet coherence between stimulus and response. While WT and 5xFAD mice exhibited age-related decline in retinal complexity, Tg^XBP1s^ and Tg^XBP1s^/5xFAD mice maintained higher complexity levels and increased Wcoh in adulthood, indicating functional preservation. These results demonstrate that sus-tained XBP1s expression protects retinal electrophysiological integrity and highlight the retina’s value as a scalable, noninvasive biomarker platform to evaluate therapeutic efficacy targeting neurodegenerative mechanisms.

## INTRODUCTION

The unfolded protein response (UPR) effector X-box binding protein 1 spliced form (XBP1s) has emerged as a potent neuroprotective modulator that restores endoplasmic reticulum homeostasis and proteostasis under stress and during aging ^1,2^. While previous work in the 5xFAD Alzheimer’s disease (AD) mouse model demonstrated that XBP1s upregulation reduces amyloid-*β* deposition in the brain, preserves synaptic function, and sustains cognition ^3–5^, its translational potential depends on the model we use for an accessible functional readout. Here, we focus on XBP1s as a therapeutic target, employing the retina as a tractable platform derived from the central nervous system (CNS) to test the consequences of XBP1s gene delivery in the 5xFAD mouse.

The retina recapitulates key anatomical, cellular, and metabolic properties of the brain–sharing embryological origin, neuronal-glial interactions, a similar vascular network, and high proteostatic demands–yet offers unparalleled accessibility for both *in vivo* and *ex vivo* assessment ^6,7^. In AD and aging, retinal thinning, microvascular alterations, and amyloid-*β* accumulation parallel cortical atrophy and cognitive decline ^8–14^. Building on evidence that XBP1s deficiency accelerates retinal degeneration ^15^, we generated double transgenic mice by crossing XBP1s-overexpressing animals (Tg^XBP1s^) with the 5xFAD AD model, yielding Tg^XBP1s^/5xFAD progeny that retain XBP1s expression from the parental Tg^XBP1s^ line despite carrying the AD-associated genes ^4^. We therefore hypothesize that XBP1s overexpression preserves retinal electrophysiological dynamics even in the context of AD pathology.

Importantly, theoretical frameworks of aging and disease increasingly point to a loss of physiological complexity, defined as a reduction in adaptive, multiscale coordination among system components, as a hallmark of dysfunction ^16–18^. Multiscale entropy (MSE) enables quantification of such complexity across temporal scale ranges ^19,20^ and has been effectively used to detect complexity loss in neurodegenerative conditions and brain injury ^21–23^.

In this study, we assessed retinal function via multielectrode array (MEA)-recorded micro-electroretinograms (*µ*ERG), analyzing advanced MSE and wavelet coherence (Wcoh) measures to capture signal complexity and stimulus-response synchronization. Our findings demonstrate that retinal electrophysiological complexity and circuit dynamics are preserved, at the ages tested, in Tg^XBP1s^/5xFAD mice. This is reflected in the recovery of *µ*ERG MSE and Wcoh metrics to near-wild-type levels, thereby effectively preventing the age-and AD-related functional decline observed in 5xFAD animals. This dual efficacy highlights XBP1s not only as a promising genetic therapy for AD but also as a key modulator of neuronal resilience in normal aging. By placing XBP1s augmentation center stage and leveraging the retina as a highly sensitive functional assay, our work establishes a clear framework for UPR-targeted interventions in both neurodegenerative and age-related contexts.

## RESULTS

### Complexity of *µ*ERG signals decays with age, but not for XBP1s protected animals

*µ*ERG signals, the light-evoked responses from photoreceptor and bipolar cells, were recorded in response to a chirp visual stimulus and a white noise (WN) checkerboard flickering (Fig. 1B). The area under the MSE curve represents an estimation of the complexity of the signal ^19^. Random signals usually considered complex, *e.g.*, pink noise (Fig. 2A), exhibit a MSE curve with consistently high values of entropy across scales. However, the entropy values can diminish markedly when a periodic component is introduced, *e.g.*, pink noise combined with a 10-Hz sine wave (Fig. 2B). Conversely, a simpler random signal like WN exhibits decreasing entropy for coarser scales (Fig. 2B). Qualitatively, the refined composite MSE (RCMSE) curves of *µ*ERG signals in response to the chirp exhibited similar shape to those of complex signals, and the average curve for each genotype, *i.e.*, wild-type (WT), Tg^XBP1s^, 5xFAD, and Tg^XBP1s^/5xFAD differed for high scales (Fig. 2C), with the WT and 5xFAD adult groups showing lower entropy for scales *>∼* 25. This resulted in significantly lower complexity for high scales–as quantified as cumulative entropy for scales from 31 to 45, as used in previous studies ^24^–for adult retinas of WT and 5xFAD subjects as compared to young retinas of the same genotypes, respectively (*p <* 0.01, multiple comparisons Tukey’s test after two-way ANOVA) (Fig. 2F). The mean (*±* standard deviation) complexity for high scales was: 0.259 (*±*0.063, *n* = 13), 0.240 (*±*0.055, *n* = 9), 0.250 (*±*0.049, *n* = 9), and 0.229 (*±*0.055, *n* = 6) for young WT, Tg^XBP1s^, 5xFAD, and Tg^XBP1s^/5xFAD, respectively, and 0.182 (*±*0.046, *n* = 10), 0.238 (*±*0.045, *n* = 10), 0.185 (*±*0.050, *n* = 11), and 0.260 (*±*0.047, *n* = 11) for adult WT, Tg^XBP1s^, 5xFAD, and Tg^XBP1s^/5xFAD, respectively. Importantly, the complexity for the young and adult animals showed no significant differences for both the Tg^XBP1s^ and Tg^XBP1s^/5xFAD groups. Moreover, the slope of the complexity indices with respect to age (in days) was negative for 5xFAD subjects but positive for Tg^XBP1s^/5xFAD crossbred animals (Fig. 2G, and Supplementary Information (SI) Appendix, Fig. S1A), indicating an age-related decline in AD that is absent in progeny protected with XBP1s. In addition, this slope was significantly different from zero (F test) for WT (*p <* 0.05) and 5xFAD (*p <* 0.01), but not for Tg^XBP1s^ (*p* = 0.555) or Tg^XBP1s^/5xFAD (*p* = 0.106). These findings suggest that XBP1s helps preserve function in the aged retina since Tg^XBP1s^/5xFAD crossbred animals–despite carrying the AD-linked transgenes–retain high complexity and avoid age-related loss, similar to non-diseased animals.

**Figure 1:**
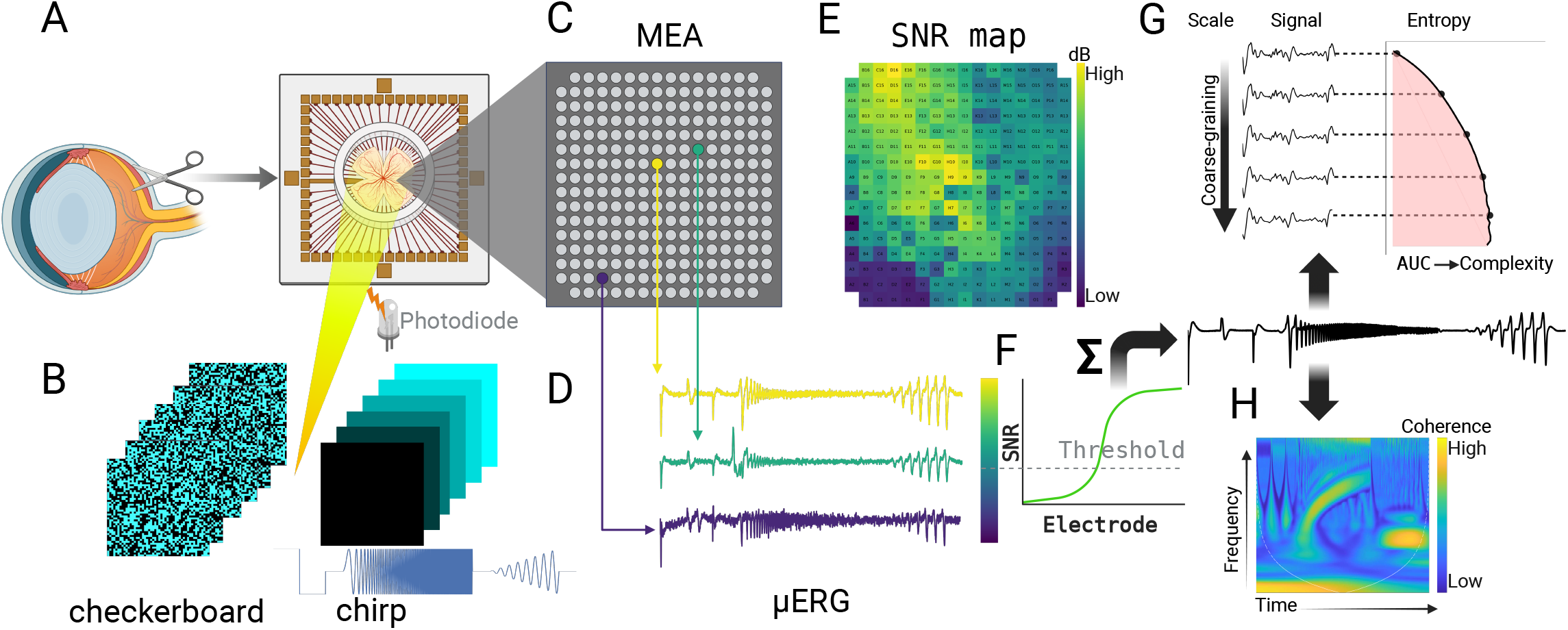
Electrophysiological recordings of retinal responses were obtained using a multielectrode array (MEA). (A) Retinal explants were isolated and placed onto a 252-MEA for *ex vivo* micro-electroretinogram (*µ*ERG) recordings. (B) Visual stimuli were delivered via a DLP projector: a white noise (WN) checkerboard pattern, and a chirp consisting of three sequential phases: an ON-OFF phase with a 3 s ON-light flash followed by 3 s of OFF-dark period; a frequency sweep phase featuring a sinusoidal light stimulus of fixed intensity with increasing frequency (1–10 Hz); and an intensity sweep phase with a 1 Hz sinusoidal stimulus of progressively increasing intensity. The actual light intensity was simultaneously recorded using a photodiode. (C) Layout of the 252-electrode MEA. (D) Raw *µ*ERG signals were band-pass filtered (0.1–40 Hz), downsampled to 250 Hz, and stored for offline analysis. (E) Electrode signal quality was assessed by computing the signal-to-noise ratio (SNR) of each channel. (F) Only electrodes with SNR *>* 7 dB were retained, and their signals were averaged for subsequent analysis. (G) Signal complexity was assessed using MSE, calculated by estimating fuzzy entropy across progressively coarse-grained versions of the signal. (H) Stimulus-response synchrony was evaluated using wavelet coherence (Wcoh) analysis based on the continuous wavelet transform (CWT) of the photodiode and *µ*ERG signals, providing a joint time-frequency representation of coherence. Created in BioRender.

**Figure 2:**
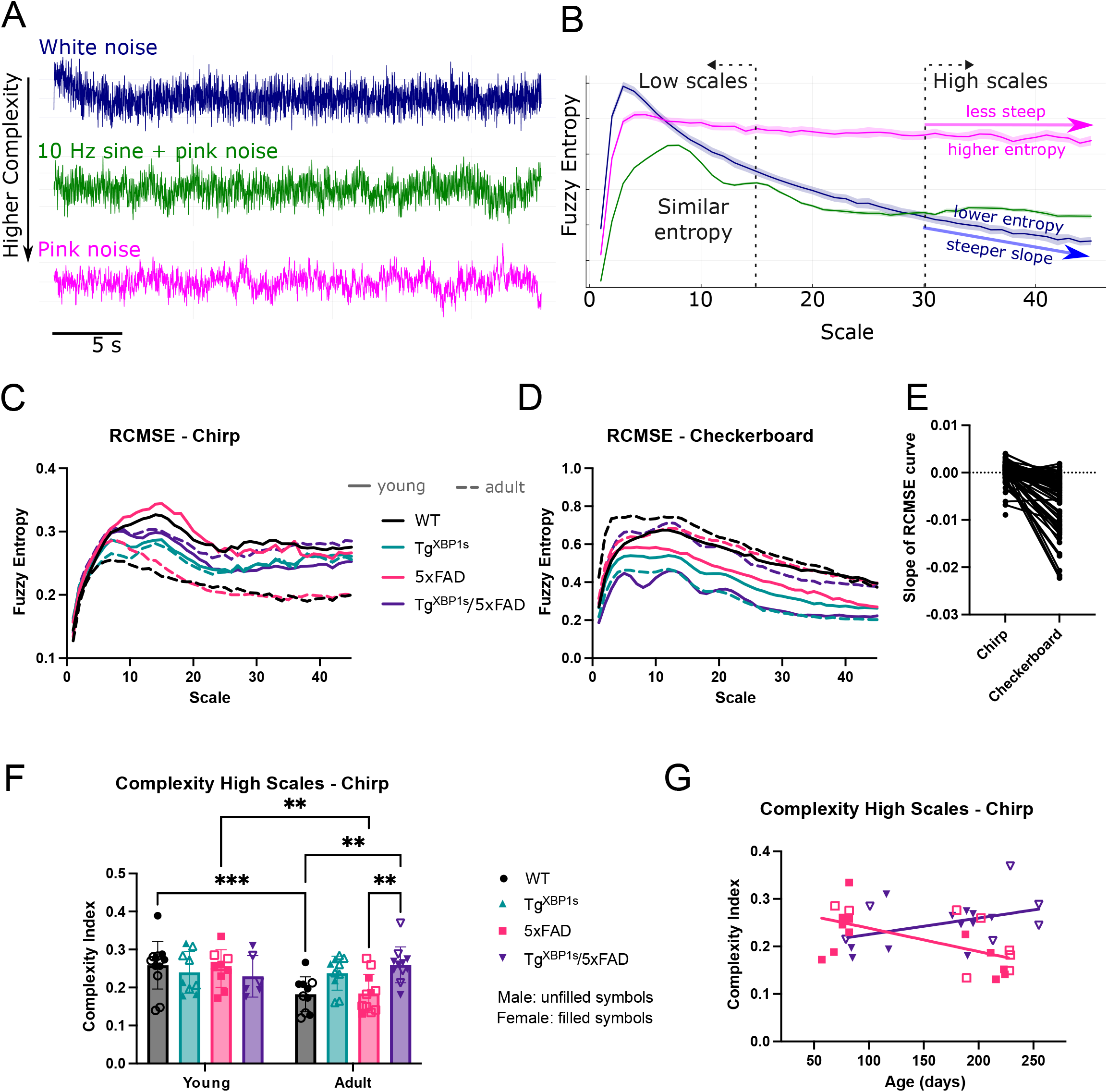
Complexity of *µ*ERG signals declines with age, but is preserved in XBP1s-overexpressing animals. (A) Sample traces of white noise (WN), a 10 Hz sinusoid plus pink noise, and pink noise (band-pass filtered at 0.1–40 Hz, sampled at 250 Hz) are shown for illustration. Pink noise is generally considered a more complex signal than WN. (B) RCMSE curves for these signals (lines: mean; shaded areas: standard deviation across 10 repetitions) show that pink noise exhibits higher entropy at larger scales and a shallower slope than WN. (C, D) RCMSE curves of *µ*ERG responses to chirp (C) and checkerboard (D) stimuli were calculated using fuzzy entropy for WT (*n* = 23), Tg^XBP1s^ (*n* = 19), 5xFAD (*n* = 20), and Tg^XBP1s^/5xFAD (*n* = 17) mice at young and adult ages. (E) The slope of the RCMSE curve at scales *>* 30 differed between the two stimuli. (F, G) For the chirp, a complexity index was calculated as the area under the RCMSE curve at high scale range (31–45) and shown for all groups (F), and for 5xFAD and Tg^XBP1s^/5xFAD as a function of age in days (G) (lines: least-squares linear regression; 1*/*slope [95% confidence intervals for slope]: *−*2033 [ *−*8.253*, −*1.587] *×* 10^−4^, and 2918 [ *−*0.823, 7.676] *×* 10^−4^ for 5xFAD, and Tg^XBP1s^/5xFAD, respectively). (**: *p <* 0.01, ***: *p <* 0.001, Tukey’s post hoc test following two-way ANOVA).

### Complexity for low scales maintains across age for all genotypes

The average RCMSE curves of *µ*ERG signals in response to the chirp were monotonously increasing and nearly identical for all groups at low scales (*<∼* 8), regardless of age (Fig. 2C). Further, the complexity for low scales–as quantified as cumulative entropy for scales from 1 to 15–showed no statistical differences for all animal groups of both ages (SI Appendix, Fig. S1B). The mean (*±* standard deviation) complexity for low scales was: 0.263 (*±*0.076, *n* = 13), 0.248 (*±*0.035, *n* = 9), 0.275 (*±*0.076, *n* = 9), and 0.257 (*±*0.036, *n* = 6) for young WT, Tg^XBP1s^, 5xFAD, and Tg^XBP1s^/5xFAD, respectively, and 0.219 (*±*0.084, *n* = 10), 0.236 (*±*0.041, *n* = 10), 0.241 (*±*0.070, *n* = 11), and 0.262 (*±*0.042, *n* = 11) for adult WT, Tg^XBP1s^, 5xFAD, and Tg^XBP1s^/5xFAD, respectively. Therefore, the age-dependent reduction of complexity observed in WT and 5xFAD retinas manifested only in coarse temporal scales.

### Random stimuli evoke responses of undistinguishable entropy content

The checkerboard-like stimulus–a realization of WN generated by sampling pixel values from a uniform random distribution–resulted in RCMSE curves that resembled that of WN (Fig. 2B,D). Further, these RCMSE curves exhibited steeper, more negative slopes than those observed for the chirp (Fig. 2E). Remarkably, although the complexity indices obtained for this stimulus were higher than those obtained for the chirp, there were no differences in complexity among the groups, regardless of the scale range (SI Appendix, Fig. S2). In sum, a purely stochastic stimulus elicited retinal responses with high entropy content that receded at coarser scales–an inherent feature of simple noise signals–thereby mirroring the entropy profile of WN.

### *µ*ERG responses are coherent in frequency and time with sinusoidal stimuli

To further characterize the *µ*ERG responses to the more meaningful chirp stimulus, we quantified the coherence between the Continuous Wavelet Transform (CWT) of these responses and the CWT of the photodiode signal recorded during the application of the chirp (Fig. 3A,B), thereby obtaining the wavelet coherence (Wcoh) as a metric of stimulus-response synchrony in the time and frequency domains (Fig. 3C). Wcoh effectively detects transient coupling between non-stationary neural signals ^25^, outperforming traditional Fourier-based methods ^26^. The *µ*ERG responses of all animals exhibited pronounced Wcoh with the chirp stimulus in frequency bands that closely reflected the chirp’s spectral composition (Fig. 3D–G). During the frequency sweep phase, Wcoh peaked in the *∼*1–10 Hz range, consistent with the increasing frequency of the stimulus (Fig. 3A.I, stimulus 2). Likewise, during the intensity sweep phase–delivered at a constant 1 Hz–Wcoh was highest around 1 Hz (Fig. 3A.I, stimulus 3). These patterns were evident across all groups in the average Wcoh maps, though specific differences emerged in defined time-frequency regions (Fig. 3D–G,a– e).

**Figure 3:**
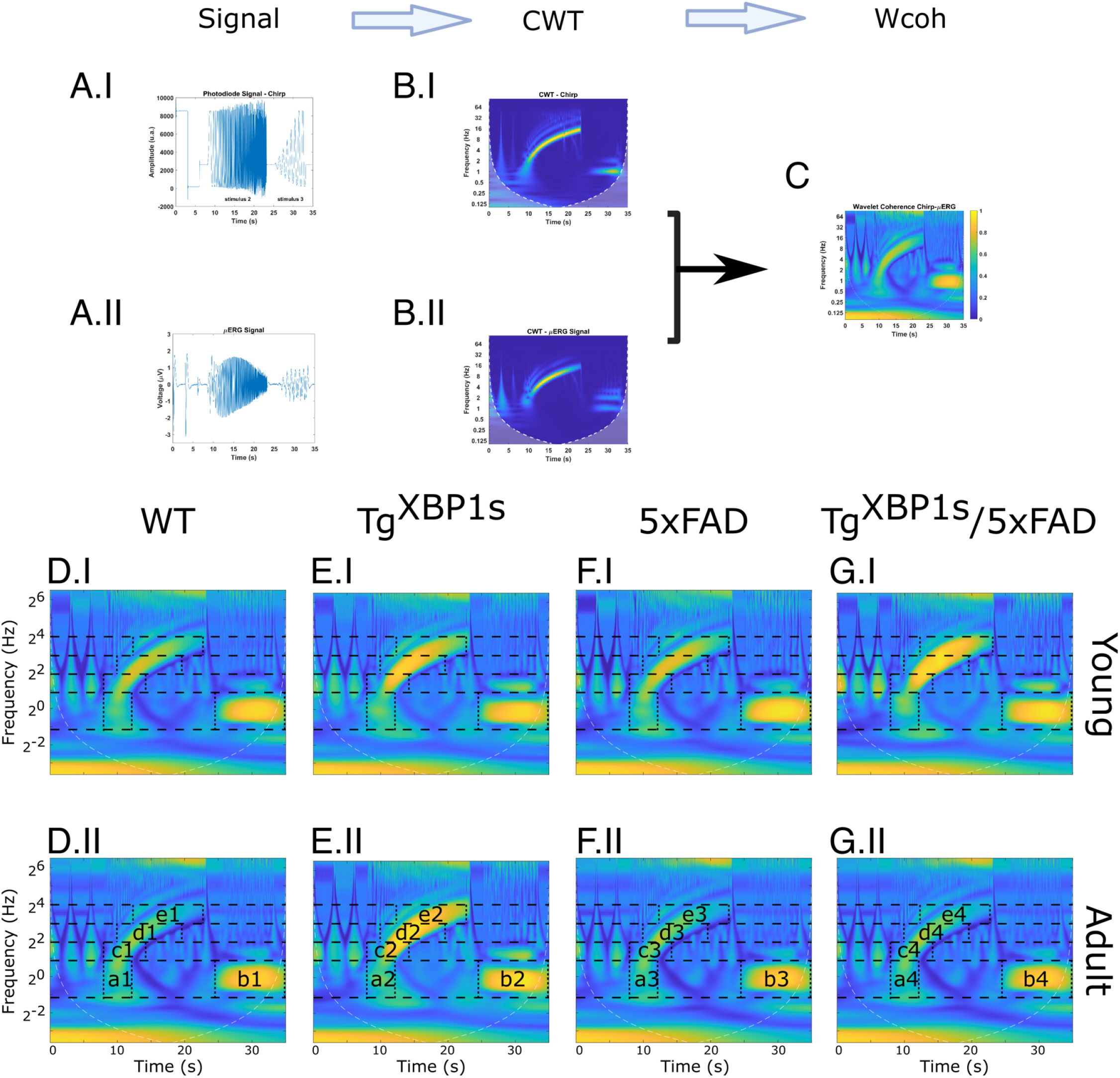
Time-frequency analysis of chirp-evoked *µ*ERG signals using continuous wavelet transform (CWT) and Wcoh. (A, B) Ten consecutive recordings of the photodiode signal (A.I) and the corresponding *µ*ERG responses (A.II) were analyzed. The frequency sweep phase (stimulus 2) and the intensity sweep phase (stimulus 3) of the chirp stimulus were considered for time-frequency decomposition. The CWT was applied to both the stimulus (B.II) and the retinal responses (B.II). (C) Wcoh was calculated for each stimulus-response pair. The resulting coherence maps were averaged across the 10 repetitions per electrode, and electrodes with SNR *>* 7 dB were included in the final average to generate a time-frequency representation of Wcoh for each animal. (D,E,F,G) Group-level Wcoh maps were obtained for each genotype and age group: WT (D), 5xFAD (E), Tg^XBP1s^ (F), and Tg^XBP1s^/5xFAD (G). Five time-frequency regions were defined for subsequent analysis in each group and model; a: [8–13 s, 0.5–2 Hz], b: [25–35 s, 0.5–2 Hz], c: [8–15 s, 2–4 Hz], d: [10–20 s, 4–8 Hz], e: [13–23 s, 8–16 Hz].

### XBP1s enhances coherence at specific frequency bands

We then performed a detailed analysis of Wcoh across specific frequency bands (4–8 Hz and 8–16 Hz) and time intervals to investigate age- and genotype-dependent modulations of stimulus-response synchrony (Fig. 4). The temporal dynamics of Wcoh (Fig. 4A,B, and SI Appendix, Fig. S3) revealed clear stimulus-evoked coherence with peaks typically occurring around 10–15 s for the 4–8 Hz band (region d), and around 15–20 s for the 8–16 Hz band (region e), for both young and adult animals across all genotypes.

**Figure 4:**
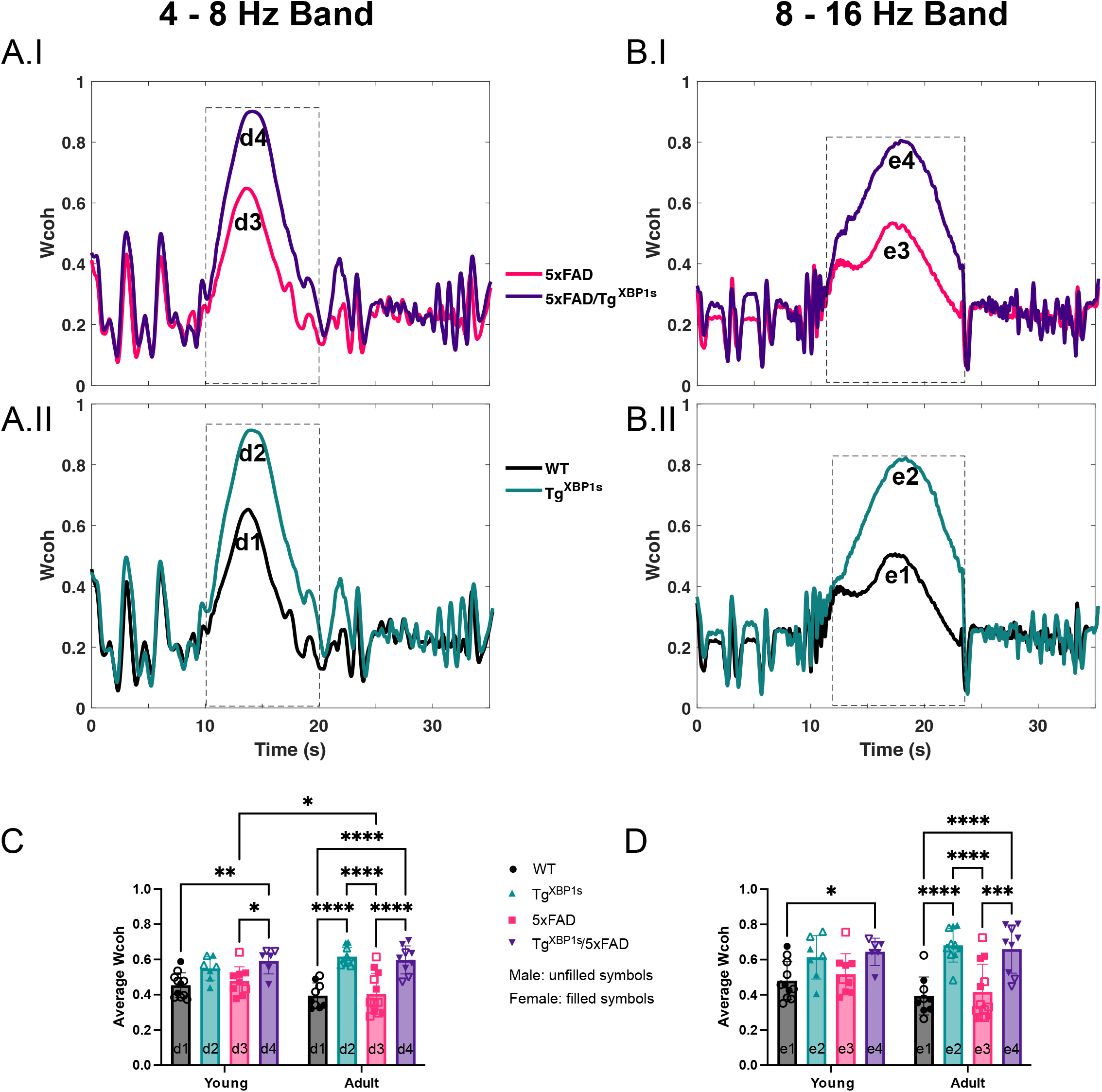
Stimulus-response synchronization quantified as Wcoh improves in XBP1s-overexpressing animals in adulthood. (A,B) Temporal evolution of Wcoh for adult animals (*n* = 9, 10, 12, 9 for WT, Tg^XBP1s^, 5xFAD, and Tg^XBP1s^/5xFAD, respectively) in the 4–8 Hz (A.I,II) and 8–16 Hz (B.I,II) bands (Wcoh for young animals, and for additional frequency bands are included in the SI Appendix). (C,D) Within selected time windows (dashed rectangles in A and B), average Wcoh amplitudes were quantified and summarized in bar plots: 4–8 Hz band (C: d1–d4), and 8–16 Hz band (D: e1–e4). (*: *p <* 0.05,**: *p <* 0.01,***: *p <* 0.001,****: *p <* 0.0001, Tukey’s post hoc test following two-way ANOVA).

Quantification of average Wcoh amplitudes within specific time windows (indicated by dashed rectangles in Fig. 4A,B and SI Appendix, Fig. S3) showed notable effects based on genotype rather than age (Figure 4C,D and SI Appendix, Fig. S4). In the 4–8 Hz band (Fig. 4C), adult Tg^XBP1s^ and Tg^XBP1s^/5xFAD mice exhibited significantly higher Wcoh values (d2, adult 0.616 *±* 0.051, *n* = 10, d4, adult 0.593 *±* 0.075, *n* = 9) as compared to age-matched 5xFAD mice (d3 adult 0.405 *±* 0.115, *n* = 12,with *p <* 0.001 in both d2–d3 and d4–d3 comparisons), suggesting that the XBP1s transgene counteracts the genotype-specific decline in coherence observed in 5xFAD animals. In particular, while WT mice did not show a significant difference between age groups (d1, young 0.454 *±* 0.070, *n* = 11, adult 0.395 *±* 0.072, *n* = 9), a significant reduction was observed in aging 5xFAD mice (d3, young 0.478 *±* 0.081, *n* = 9, adult 0.405 *±* 0.115, *n* = 12, *p <* 0.05). In contrast, coherence in Tg^XBP1s^ and Tg^XBP1s^/5xFAD mice remained high and stable across age groups, suggesting a XBP1-mediated preservation of stimulus-response synchrony despite aging (d2 young 0.549 *±* 0.066, *n* = 7, d4 young 0.597 *±* 0.081, *n* = 6).

Similar patterns were observed in the 8–16 Hz band (Fig. 4D), albeit with differences more centered within the adult animals. More specifically, both the Tg^XBP1s^ and Tg^XBP1s^/5xFAD adult group exhibited significantly higher Wcoh than age-matched WT and 5xFAD mice (e1–e2: *p <* 0.0001, e2–e3: *p <* 0.0001, e1–e4: *p <* 0.001, e3–e4: *p <* 0.001; with e1: 0.393 *±* 0.108, *n* = 9, e2: 0.681 *±* 0.095, *n* = 10, e3: 0.416 *±* 0.158, *n* = 12, e4: 0.651 *±* 0.130, *n* = 9), reinforcing the protective effect of XBP1s on stimulus-response synchrony.

Although Wcoh did not show marked effects based on age for both frequency bands, it did show a downward trend in the WT and 5xFAD groups, as indicated by the negative slopes of Wcoh vs age in days (SI Appendix, Fig. S5).

Collectively, these results demonstrate that while stimulus-response synchrony, as measured by Wcoh, declines with age in control and 5xFAD models, the presence of the XBP1s is associated with its remarkable preservation in adult mice across relevant frequency bands.

## DISCUSSION

Building on the established neuroprotective role of XBP1s in the brain, we hypothesized that its upregulation in the retina may similarly mitigate age-related degeneration and preserve visual function. In WT and 5xFAD mice, we observed a progressive decline in complexity and Wcoh–albeit more modest–with age (Figs. 2G, and SI Appendix, Fig. S5). However, Tg^XBP1s^ and Tg^XBP1s^/5xFAD animals maintained higher levels for both metrics, suggesting that XBP1s expression confers functional resilience to retinal circuits.

AD pathology begins well before the onset of clinical symptoms ^27^. While biomarkers such as tau and amyloid-*β* are clinically informative, current detection methods–including cerebrospinal fluid analysis and PET imaging–are invasive, expensive, and not broadly scalable. In contrast, the retina’s noninvasive accessibility creates a powerful translational window for detecting and monitoring neurodegenerative processes. In both patients and animal models, AD-related pathology produces characteristic alterations in ERG waveforms, namely, reduced amplitudes and delayed latencies ^24,28–32^. Although our findings did not reveal significant differences in WT and 5xFAD animals, we observed robust age-related changes in *µ*ERG complexity. By applying RCMSE and Wcoh analyses to *µ*ERG recordings, our study provides a more sensitive and quantitative readout of neural integrity and therapeutic efficacy. This approach supports the retina as an accessible, cost-effective platform for preclinical evaluation of neuroprotective interventions. In our data, aging in WT and 5xFAD mice resulted in reduced complexity, particularly at higher scales associated with integrative retinal processes. Conversely, XBP1s-expressing mice preserved signal complexity with age, suggesting maintenance of hierarchical and coordinated retinal dynamics. Further, for these animals, Wcoh analysis showed enhanced stimulus-response coherence in adulthood in mid-frequency bands (4–8 Hz and 8–16 Hz), likely reflecting preserved inner retinal integration (photoreceptors, horizontal, bipolar and amacrine cells) synchrony. Importantly, genotype-dependent differences in entropy emerged only at higher scales, while lower-scale entropy, corresponding to fast, local computations, remained stable across all groups. This suggests that aging primarily disrupt large-scale, integrative processes while sparing local circuit activity, echoing findings from cortical systems ^33,34^.

The nature of the applied stimulus was also critical in revealing these effects. Structured chirp stimuli, which modulate frequency and intensity over time, reliably differentiated between experimental groups. In contrast, the checkerboard WN stimulus, composed of temporally random patterns, produced responses with entropy profiles similar across all genotypes, likely due to nonspecific, stochastic retinal activation (Fig. 2B,D). These results underscore the importance of structured, *e.g.* chirp-like, stimuli in probing the integrity of temporal processing and complexity in sensory systems. Moreover, alternative multidimensional entropy metrics, such as permutation entropy ^35^, interscale dissimilarity ^36^, or spatially resolved 2D/3D entropy, may provide even greater sensitivity to detect dysfunction, particularly under noisy or unstructured input conditions.

Gender is a recognized confounding factor in AD, and sex differences in both behavioral and molecular pathology have been reported in 5xFAD mice ^37^. In this study, female 5xFAD mice exhibited lower, albeit non-significant, *µ*ERG complexity compared to males in both the young group (0.241 *±* 0.049 vs. 0.281 *±* 0.047, *p* = 0.356, Tukeys test following two-way ANOVA) and the adult group (0.167 *±* 0.035 vs. 0.199 *±* 0.053, *p* = 0.892, Tukeys test following two-way ANOVA). However, the number of males in the young group was smaller, whereas the sex distribution was more balanced in the adult group. Consequently, these observed differences in complexity may have been influenced by disparities in group sizes between sexes. Future studies should include larger and more balanced cohorts to systematically investigate sex differences in MSE metrics of the *µ*ERG in both male and female mice.

While we did not perform molecular analyses of the retina, prior work in the hippocampus of 5xFAD mice demonstrated that XBP1s overexpression (via crossbreading) restores up to 76% of disease-associated proteomic disruptions–including deficits in synaptic function, cytoskeletal integrity, and ER proteostasis ^4^. Retinal molecular and histochemical studies remain to be conducted, but existing literature supports XBP1s’s role in mitigating ER stress, preserving glycolytic metabolism, and maintaining synaptic integrity ^4,15^. Together, these findings establish XBP1s as a pivotal genetic modulator of retinal aging and degeneration, reinforcing the retina’s utility as a sensitive, accessible platform for neurodegenerative research and therapeutic evaluation ^17,38^. Our work thus underscores the translational promise of targeting XBP1s to preserve neural health and to develop noninvasive retinal biomarkers for early detection and monitoring of neurodegenerative diseases.

## MATERIALS AND METHODS

Methods are described in detail in SI Appendix.

### Animals

All experimental protocols adhered to the bioethical guidelines established by the Universidad de Valparaíso, aligned with international standards for the care and handling of animals (protocol No. BEA15620), and the bioethics and biosecurity standards of the Chilean National Agency for Research and Development (ANID). We used four animal groups, each one divided in two age groups: WT mice (*n* = 13 young, *n* = 10 adult), Tg^XBP1s^ transgenic mice (*n* = 9 young, *n* = 10 adult) engineered to overexpress XBP1s, 5xFAD mice (*n* = 9 young, *n* = 11 adult) as a robust model of AD, and Tg^XBP1s^/5xFAD mice (*n* = 6 young, *n* = 11 adult), the crossbreed of Tg^XBP1s^ and 5xFAD animals ^4^. The gender and age characteristics are summarized in Table S1.

### Electroretinography (*µ*ERG)

Electrophysiological function of isolated retinal tissue was assessed using 252-electrode MEA for *µ*ERG recordings, including principally the response from photoreceptors and bipolar cells, as previously described ^39,40^. Visual stimuli were delivered via a calibrated DLP projector equipped with specialized optics to focus a 400 *×* 400 pixel image onto the MEA. Given the dichromatic vision of mice, we used cyan stimuli (blue + green) as basal stimulation, calibrated with peak emissions at 460 nm and 520 nm. The stimulation protocol included a chirp and a WN stimuli. The chirp had three phases: (1) an 3-s each ON and OFF flash; (2) a frequency sweep consisting of a 15-s sinusoidal stimulus increasing from 1 to 15 Hz; and (3) an intensity sweep comprising an 8-s sequence of eight incremental intensity steps. The WN was a sequence of cyan checkerboard patterns (0.05 mm block size, 35Œ35 grid) at 60 Hz.

Mean irradiance at the MEA surface was calibrated at 70 nW/mm^2^. Recordings were filtered using a band-pass filter (0.1-40 Hz), resampled to 250 Hz, normalized with z-score, and averaged across 10 trials to obtain a representative response to a stimulus per electrode. Finally, channels were selected according to a signal-to-noise (SNR) threshold criterion of 7 dB, so that only selected channels were averaged for further analysis (median number of selected channels [min–max]: 145 [1–252]).

### Multiscale entropy and complexity

We used the concept of entropy to quantify the irregularity of *µ*ERG signals. More specifically, we calculated the Fuzzy Entropy (FuzzyEn) ^41^ of the signals, which is an extension of the so called sample entropy. To capture signal dynamics across multiple time scales, we applied the RCMSE method ^42^, which calculates the entropy over coarse-grained versions of the original signal using a refined averaging strategy. For a given scale *τ*, coarse-grained time series are constructed by averaging *τ* consecutive, non-overlapping points starting at different offsets. Entropy is computed for each series and the resulting values are averaged before applying the logarithm, improving statistical reliability and reducing the risk of undefined values at larger scales. FuzzyEn was computed using parameters *m* = 2 and *r* = 0.2, up to scale 45, based on a minimum of 50 samples per scale. We calculated complexity indices as the normalized area under the curve calculated over a specified scale range ^20^.

### Wavelet coherence

We calculated the Wcoh between the chirp stimulus, *i.e.*, recorded photo-diode signal, and the corresponding *µ*ERG response for 10 stimulus repetitions at each electrode. The Wcoh average of these repetitions and of selected electrodes was obtained for each subject. Wcoh was defined as the time-frequency representation of the normalized cross-wavelet transform of the two signals. We then determined the time course of Wcoh in each band of interest and calculated the weighted area under the curve as a measure of each animal’s stimulus-response coherence in that frequency band ^25^. Signal processing and corresponding graphs were obtained using Matlab software.

### Statistical analysis

For the complexity index and Wcoh, we performed Shapiro-Wilk test of normality. Then, we performed two-way analysis of variance (ANOVA) with age and genotype as factors, followed by multiple comparisons Tukey’s test. For data arranged by age in days, we applied least-squares linear regression followed by an F-test to determine if the slope was significantly non-zero. In all cases, we set a significance level *α* of 0.05. All statistical analyses were performed in GraphPad Prism v10.

## ACKNOWLEDGMENTS

Supported by the Chilean National Agency for Research and Development (ANID) through Grant Exploración No. 13220082 to LEM and AGP, Fondecyt No. 1200880 and Fondequip No. EQM190032 to AGP, and Basal No. FB210017 to CR. Computational support as cluster access was provided by Universidad de Santiago de Chile; we thank Cristóbal Acosta for technical assistance in the use of this cluster.

## AUTHOR CONTRIBUTIONS

LEM, and AGP conceived, supervised, and obtained funding for the study. JAA, DN, and DP performed the *ex vivo* experiments. JB, MC, CR, LEM, and AGP analyzed the data. MO, BC, JB, FM, and LEM implemented the computational methods. CH and CDA engineered the Tg^XBP1s^ animals and commented on the text. JB, CR, LEM, and AGP wrote the manuscript.

