## Supplementary Information Appendix for "XBP1s as a Therapeutic Target to Preserve Retinal Function During Aging and Neurodegeneration"

### MATERIALS AND METHODS

#### Electroretinography

Mice were subjected to a 12-12 light/dark cycle in a temperature-controlled environment, with *ad-libitum* access to water and food. Before each experiment, the animals were dark-adapted for 30 minutes, then deeply anesthetized with isoflurane (Baxter, Deerfield, IL, USA) and sacrificed. The eyes were quickly enucleated under dim red light and the eyecups were immersed in Ames medium with bicarbonate buffer (Sigma-Aldrich, St. Louis, MO, USA) at 32 řC and pH 7.4, continuously oxygenated with 95% ($O_2$) and 5% ($CO_2$). Small pieces of retina were gently separated from the retinal epithelium and placed on a rod device that supports a dialysis membrane ring (MWCO-25000, Spectrumlabs, Rancho Dominguez, CA, USA), treated with polylysine (Product P4707, Sigma-Aldrich, St. Louis, MO, USA) to facilitate contact between the retina and the surface of the MEA electrodes.

Visual stimuli were generated using custom MATLAB scripts and delivered via a calibrated DLP projector equipped with specialized optics to focus a 400 Œ 400 pixel image–each pixel approximately 5 $\mu$m in diameter–onto the MEA array. Retinal samples were positioned on the array and viewed for control under an inverted microscope (Eclipse T200, Nikon). The stimulation protocol included a chirp stimulus with three phases: (1) an 3-s each ON and OFF flash; (2) a frequency sweep consisting of a 15-s sinusoidal stimulus increasing from 1 to 15 Hz; and (3) an intensity sweep comprising an 8-s sequence of eight incremental intensity steps. Chirp sequences were repeated over 10 or 21 trials depending on the experiment. Mean irradiance at the MEA surface was calibrated at 70 nW/mm$^2$. Given the dichromatic vision of mice, we used a cyan stimuli (blue + green), calibrated with peak emissions at 460 nm and 520 nm using a USB4000 Fiber Optic Spectrometer (Ocean Optics). In addition, we presented a sequence of WN cyan checkerboard patterns (0.05 mm block size, 35 X 35 grid) at 60 Hz for 30 minutes. All electrophysiological recordings were stored for offline analysis.

#### Signal pre-processing

We filtered the recordings from each electrode or channel using a band-pass filter (0.1-40 Hz), and then resampled each channel to 250 Hz. Next, we normalized the signals using z-score measurement, and averaged 10 trials to obtain a representative response to a stimulus per electrode. Finally, we selected channels according to a signal-to-noise (SNR) criterion, so that only selected channels were averaged for further analysis. For SNR estimation, the power ratio between background noise and the $\mu$ERG signal was calculated in decibels (dB), with signal power obtained for the 35 s of the chirp presentation, and the background noise power, for the immediate 4 s after the chirp presentation. We applied an SNR threshold of 7 dB.

#### Multiscale entropy and complexity

SampEn measures the unpredictability of a time series, so that higher values indicate greater level of disorder. For a time series $X$ of length $N$, and sequences of length $m$, SampEn is defined as:

$$\mathrm{SampEn}(N, m, r) = \ln \frac{\sum_{i=1}^{N-m} U_i^m(r)}{\sum_{i=1}^{N-m} U_i^{m+1}(r)} \tag{1}$$

where $U_i^m(r)$ counts the number of sequences within a distance $r$ of the $i$-th sequence, excluding self-matches. While SampEn uses a Heaviside function as a similarity metric for the distance, FuzzyEn uses a fuzzy degree of similarity, $D_{ij}^m$, which instead of being binary, depends on the distance $d_{ij}^m$ between sequences through a continuous, concave function, typically quadratic exponential:

$$D_{ij}^m = \exp(-(d_{ij}^m/r))^2. \tag{2}$$

For multiscale entropy analyses, we used the RCMSE, which improves robustness by calculating FuzzyEn for each of the $\tau$ coarse-grained time series obtained from different starting points, and averaging the resulting similarity counts before applying the logarithm. This procedure reduces the risk of undefined entropy values, and is described in detail elsewhere[42]. FuzzyEn was computed using parameters $m = 2$ and $r = 0.2$, up to scale 45, based on a minimum of 50 samples per scale.

A recently introduced theory of complexity-loss postulates that biological complexity decays with aging and disease[18,19]. In this case, biological complexity refers to the nonlinear interactions between various structural units and their collective behavior, operating over a wide range of temporal and spatial scales. Such complexity can be estimated by means of the entropy of a physiological output calculated at multiple time scales[20,43], *i.e.*, the RCMSE curve. It follows that the area under the resulting RCMSE curve can be used as an indicator of complexity of a physiological system. We calculated a complexity index as a normalized area-under-the-curve (nAUC), calculated over a specified scale range. Algorithms used in this analysis were implemented using `Julia` v1.9.3.

#### Continuous Wavelet Transform (CWT)

The CWT is performed by convolution of the analyzed function $x(t)$ with the wavelet function $\psi_{a,b}(t)$:

$$Cx(a, b) = \int_{-\infty}^{+\infty} x(t)\psi_{a,b}^*(t)dt \tag{3}$$

The function $\psi_{a,b}(t)$ (* indicates the complex conjugate) is obtained from the mother wavelet $\psi_o(t)$ (Morlet in this case):

$$\psi_{a,b}(t) = \frac{1}{\sqrt{a}}\psi_o\left(\frac{t-b}{a}\right) \tag{4}$$

The parameters $a$ and $b$ define the width and the location of the $\psi_o(t)$, and the factor $\frac{1}{\sqrt{a}}$ provides the constant unit norm of the wavelets[44].

The obtained wavelet coefficients, $Cx(a, b)$, reflect the spectral dynamics of the signal $x(t)$, since the dilation or contraction given by the scaling factor $a$ of the mother wavelet can be associated with the presence of low or high frequencies, respectively; and the translation factor $b$, with the moment at which that frequency occurs.

#### Wavelet coherence (Wcoh) analysis

Briefly, given two time series $x(t)$ and $y(t)$ with continuous wavelet transforms $C_x(a,b)$ and $C_y(a,b)$, respectively, the expression $C_{xy}(a,b) = C_x(a,b) \cdot C_y^*(a,b)$ is defined as the cross-wavelet transform, and reflects the time and frequency band in which they present a common power, including their relative phase. $|C_{xy}(a,b)|^2$ represents the power spectrum of the cross-wavelet transform, and describes the local covariance between the signals. Since high-power values of a single signal can indicate a high covariance between the two, it is common to normalize the above expression. Thus, WC is defined as:

$$\mathrm{Wcoh} = \frac{|S(C_x(a,b)C_y^*(a,b))|^2}{|S(C_x(a,b))|^2 \cdot |S(C_y(a,b))|^2} \tag{5}$$

where Wcoh is a time-frequency representation of the wavelet coherence between $x(t)$ and $y(t)$ and $S$ corresponds to a smoothing operator in time $(b)$ and scale $(a)$[45].

We calculated the Wcoh between the chirp stimulus (recorded photo-diode signal) and the corresponding $\mu$ERG response for $10$ stimulus repetitions at each electrode. The Wcoh average of these repetitions and of all the electrodes ($\mathrm{SNR} > 7$ dB) was obtained for each animal. We then determined the time course of Wcoh in each band of interest and calculated the weighted area under the curve as a measure of that animal's stimulus-response coherence in that frequency band[25]. Signal processing and corresponding graphs were obtained using Matlab software.

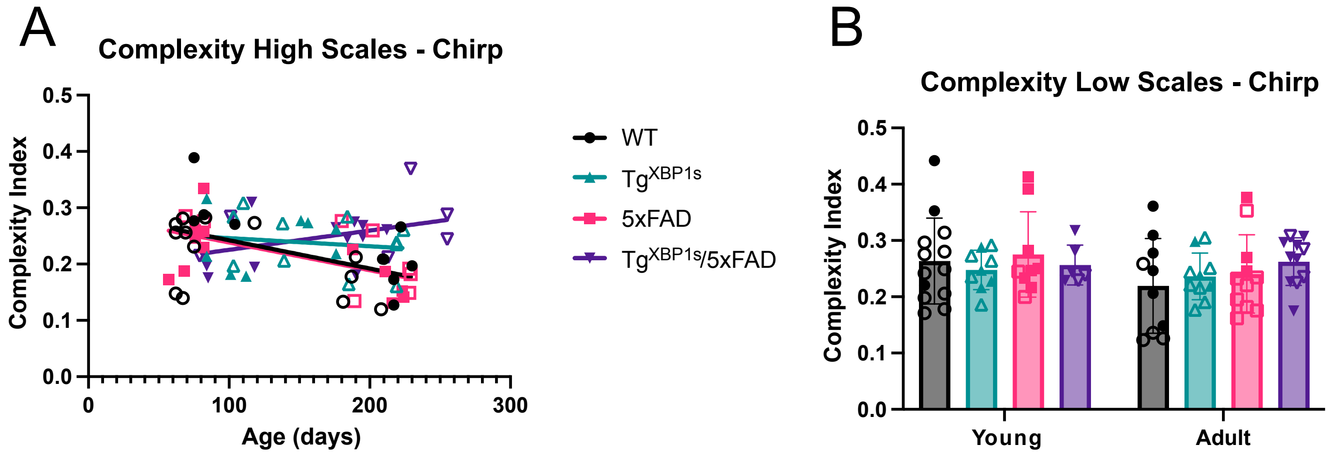


**Figure S1: Complexity of $\mu$ERG responses to the chirp stimulus decays for non-protected XBP1s animals, but only for high-scale ranges.** (A) Complexity index for high-scale (31–45) ranges as a function of age in days for WT ($n = 23$), Tg$^{XBP1s}$ ($n = 19$), 5xFAD ($n = 20$), and Tg$^{XBP1s}$/5xFAD animals ($n = 17$). Solid lines: least-squares linear regression. The corresponding values of $1$/slope [95% confidence intervals for the slope] were: $-1994$ $[-8.876, -1.153] \times 10^{-4}$, $-6917$ $[-6.508, 3.617] \times 10^{-4}$, $-2033$ $[-8.253, -1.587] \times 10^{-4}$, and $2918$ $[-0.823, 7.676] \times 10^{-4}$ for WT, Tg$^{XBP1s}$, 5xFAD, and Tg$^{XBP1s}$/5xFAD, respectively. Slope significantly different from zero (F test) for WT ($p < 0.05$) and 5xFAD ($p < 0.01$), but not for Tg$^{XBP1s}$ ($p = 0.555$) or Tg$^{XBP1s}$/5xFAD ($p = 0.106$). (B) Complexity index for low-scale (1–15) ranges for the same animals. In this case, two-way ANOVA revealed no significant interaction between genotype and age ($p = 0.617$), and there were no significant main effects of age ($p = 0.146$) or genotype ($p = 0.703$) on complexity.

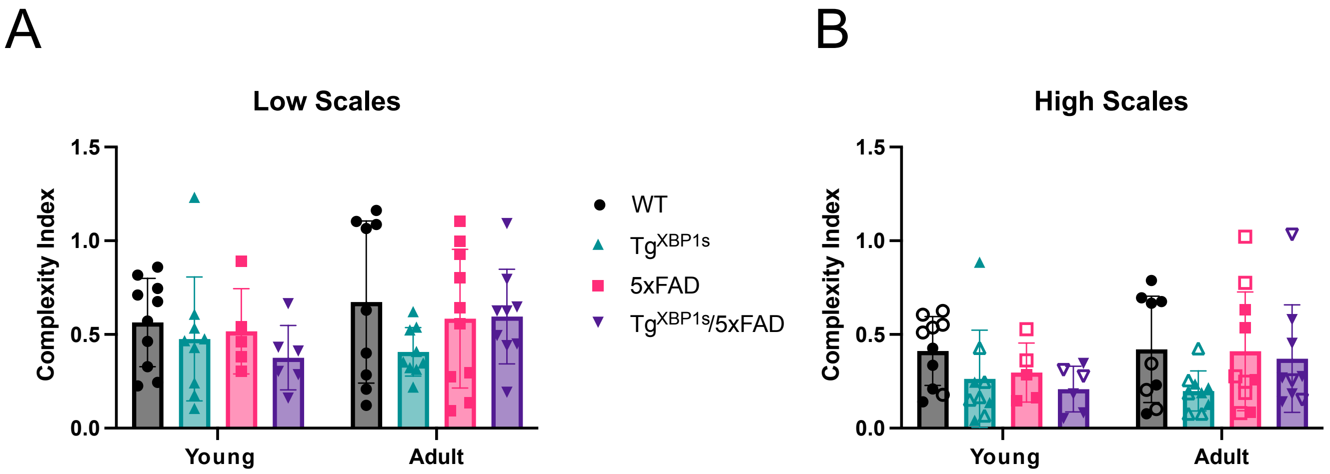


**Figure S2: Complexity of $\mu$ERG responses to the a white noise stimulus is similar for all groups.** Complexity index for low-scale (1–15) ranges (A) and high-scale ranges (31–45) ranges (B) for the checkerboard stimulus for WT ($n = 23$), Tg$^{XBP1s}$ ($n = 19$), 5xFAD ($n = 20$), and Tg$^{XBP1s}$/5xFAD animals ($n = 17$). In both cases, two-way ANOVA revealed no significant interaction between genotype and age ($p = 0.574$ and $p = 0.530$), and there were no significant main effects of age ($p = 0.272$ and $p = 0.360$) or genotype ($p = 0.300$ and $p = 0.116$) on complexity.

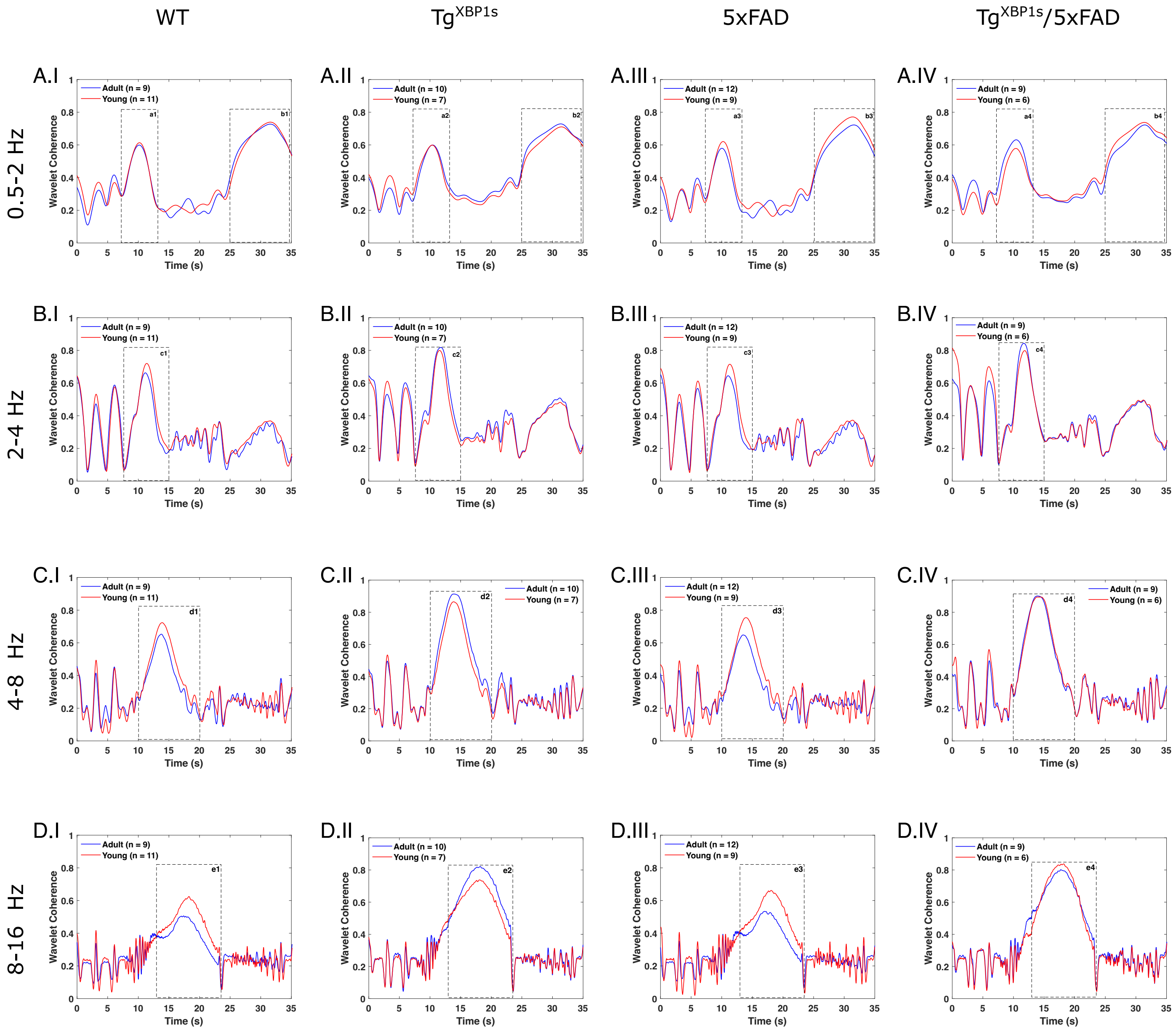


**Figure S3: Temporal evolution of Wcoh for all groups.** Wcoh was computed for each animal and averaged by group, young (solid red line) and adult (solid blue line), within each genotype (WT, $Tg^{XBP1s}$, 5xFAD, and $Tg^{XBP1s}$/5xFAD). Average traces are shown for the 0.5–2 Hz (A.I–IV), 2–4 Hz (B.I–IV), 4–8 Hz (C.I–IV), and 8–16 Hz (B.I–IV) bands. Within selected time windows (dashed rectangles), average Wcoh amplitudes were quantified for subsequent analysis: a1–a4 and b1–b4 for 0.5–2 Hz, c1–c4 for 2–4 Hz, d1–d4 for 4–8 Hz, and e1–e4 for 8–16 Hz.

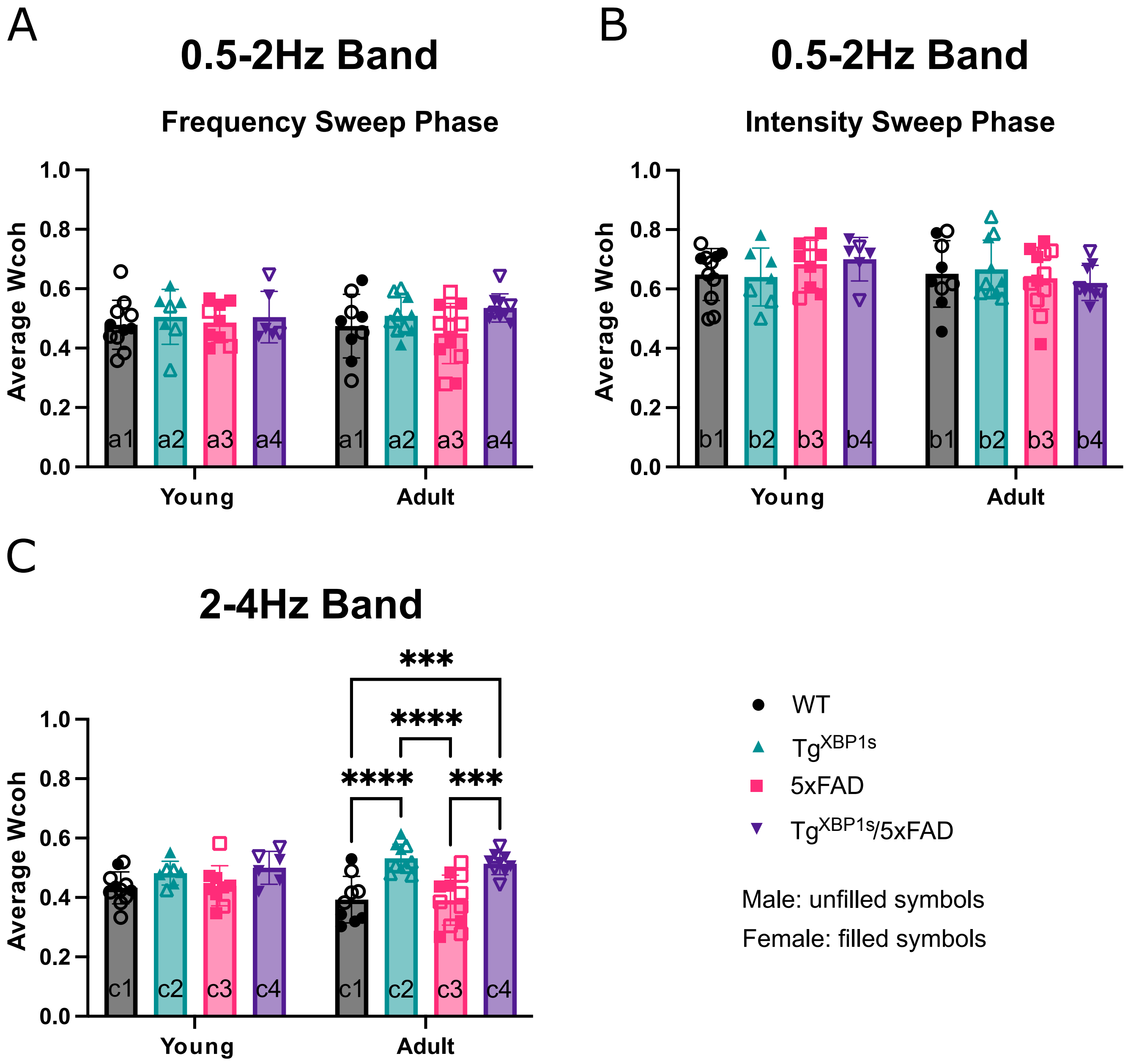


**Figure S4: Average Wcoh in very low frequency bands is comparable across all genotypes and age groups**. (A,B) Mean Wcoh in the 0.5–2 Hz band during (A) region *a*, corresponding to the frequency-sweep phase of the chirp, and (B) region *b*, corresponding to the intensity-sweep phase of the chirp. (C) Mean Wcoh in the 2–4 Hz band during region *c*. (***: $p < 0.001$, ****: $p < 0.0001$, Tukey's post hoc test following two-way ANOVA)

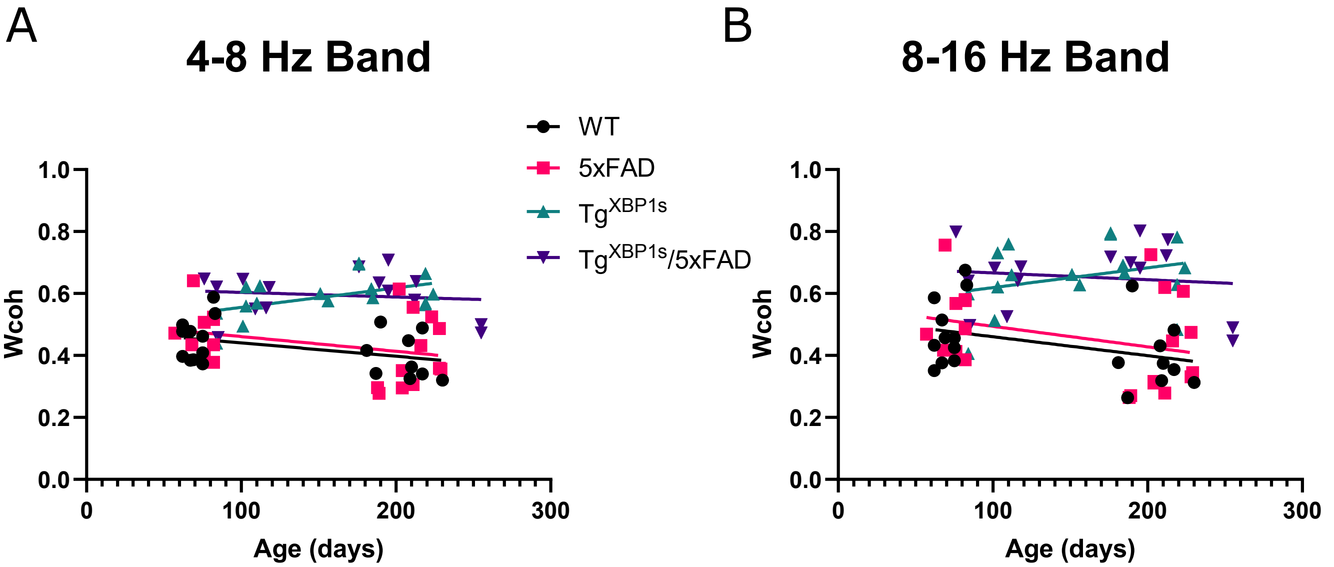


**Figure S5: Wcoh only moderately decayed with age for all groups.** Wcoh in the 4–8 Hz band (A) and in the 8–16 Hz band (B) as a function of age in days for the chirp stimulus. Solid lines: least-squares linear regression for each group. For the 4–8 Hz band, the corresponding values of $1/\text{slope}$ [95% confidence intervals for the slope] were: $-2310\ [-9.239, 0.582] \times 10^{-4}$, $1593\ [0.158, 12.40] \times 10^{-4}$, $-2110\ [-1.164, 2.162] \times 10^{-4}$, and $-6725\ [-8.548, 5.574] \times 10^{-4}$ for WT, $Tg^{XBP1s}$, 5xFAD, and $Tg^{XBP1s}$/5xFAD, respectively. For the 8–16 Hz band, the corresponding values of $1/\text{slope}$ [95% confidence intervals for the slope] were: $-1638\ [-13.61, 1.399] \times 10^{-4}$, $1571\ [-4.842, 17.57] \times 10^{-4}$, $-1513\ [-16.21, 2.989] \times 10^{-4}$, and $-4554\ [-12.98, 8.59] \times 10^{-4}$ for WT, $Tg^{XBP1s}$, 5xFAD, and $Tg^{XBP1s}$/5xFAD, respectively.

**Table 1:** Gender and age characteristics of animals.

| Genotype | Age group | $n$ | Median (min–max) euthanasia age in days | No females |
|---|---|---|---|---|
| WT | young | 13 | 75 (62–118) | 4 |
| | adult | 10 | 210 (181–230) | 6 |
| $Tg^{XBP1s}$ | young | 9 | 103 (84–139) | 4 |
| | adult | 10 | 185 (151–224) | 4 |
| 5xFAD | young | 9 | 76 (57–82) | 7 |
| | adult | 11 | 216 (180–229) | 5 |
| $Tg^{XBP1s}$/5xFAD | young | 6 | 93 (79–118) | 5 |
| | adult | 11 | 195 (176–255) | 6 |
